# Quantitative reverse transcription PCR assay to detect pyrethroid resistance in *Culex* mosquitoes

**DOI:** 10.1101/2021.05.18.444631

**Authors:** Kelli M. Hager, Erick Gaona, Amy Kistler, Kalani Ratnasiri, Hanna Retallack, Miguel Barretto, Sarah S. Wheeler, Christopher M. Hoover, Eric J. Haas-Stapleton

## Abstract

Pyrethroid insecticides are widely used to control mosquitoes that transmit diseases such as West Nile virus (WNV) to humans. A single nucleotide polymorphism (SNP) in the knockdown resistance locus (*kdr*) of the *voltage gated sodium channel* (*Vgsc*) gene of *Culex* mosquitoes confers knockdown resistance to pyrethroids. PCR-based assays that detect these SNPs in *Culex* species are currently available for *Culex pipiens* Linnaeus and *Culex quinquefasciatus* Say. RNAseq was employed to sequence the coding region of *Vgsc* for *Culex tarsalis* Coquillett and *Culex erythrothorax* Dyar, two WNV vectors. We utilized the cDNA sequence to develop a quantitative reverse transcriptase PCR assay that detects the L1014F mutation in the *kdr* of V*gsc*. Because this locus is conserved, the assay successfully detected the SNPs in multiple *Culex spp*. vectors of WNV in the United States. The resulting *Culex* RT*kdr* assay was validated using quantitative PCR, CDC bottle bioassays, and sequencing of PCR products. Using sequencing, we determined the accuracy of the *Culex* RT*kdr* assay was 99%. Pyrethroid resistance was more common among *Cx. pipiens* than other *Culex* spp. and co-occured with agriculture. We anticipate that public health and vector control agencies may utilize the *Culex* RT*kdr* assay to map the distribution of pyrethroid resistance in *Culex* species to more efficiently control mosquitoes and the diseases they transmit.

## Introduction

Many mosquitoes within the *Culex* genus that are present in California are vectors for diseases caused by West Nile virus (WNV), St. Louis Encephalitis virus (SLEV), and filarial worms [1]. WNV and SLEV are maintained in a bird-mosquito cycle by mosquitoes such as *Culex pipiens* Linneaus and *Culex erythrothorax* Dyar that preferentially feed on birds. *Culex tarsalis* Coquillett, another WNV vector, transition seasonally from ornithophilic to general feeders or when host availability is constrained [2, 3]. Humans and horses are considered dead-end hosts for these arboviruses because they generate low viremia, thereby preventing onward transmission of these arboviruses [4, 5]. There have been over 6,700 symptomatic human infections of WNV since it was introduced to California in 2003 [6, 7]. Vector control agencies interrupt disease transmission through environmental manipulation, biological or chemical control of adult and juvenile mosquitoes, and public education. Adulticides (pesticides that target biting adult mosquitoes) are used to reduce mosquito abundance and the transmission of pathogens.

Pyrethroid adulticides preferentially bind to open voltage gated sodium channels (*Vgsc*) in neuronal membranes, preventing their closure. The open *Vgsc* leaves the membrane depolarized and the neuron unable to transmit signals among cells, resulting in paralysis (i.e. knockdown) and death of the insect [8, 9]. More than 50 mutations in the sodium channel gene are associated with pyrethroid resistance among arthropods [10]. The most common among *Culex* species is the L1014F single nucleotide polymorphism (SNP), which promotes closed state inactivation and knockdown resistance [11, 12].

Pyrethroids are commonly used to control structural and agricultural arthropod pests. The CDC considers mosquito populations resistant to an adulticide when knockdown or mortality rates are less than 80% in an adult mosquito bottle bioassay (BBA; [13]). Increased use of pyrethroids in agricultural settings may contribute to pyrethroid resistance among a broad range of arthropods [14] [15]. Concerns with widespread pyrethroid resistance in mosquitoes prompted us to develop a quantitative reverse transcriptase-PCR (qRT-PCR) assay that detects the L1014F SNP in *Culex* species. Our original goal was to develop this assay for use in *Culex tarsalis*, but after comparing the cDNA sequences of other *Culex* vectors we discovered the qRT-PCR assay produced a more conserved template compared to its qPCR counterparts. Here we describe the development of a *Culex* RT*kdr* assay and an application of the assay to map pyrethroid resistance within Alameda County (California, USA).

## Methods

### 1. Mosquito Collection

Adult mosquitoes used for RT*kdr* testing were collected overnight from May - October of 2019 in Alameda County (California, USA) using Encephalitis Vector Survey traps (BioQuip, Rancho Dominguez, CA) that were baited with dry ice. A scientific collection permit was not required because the collections were made by a mosquito abatement district that was operating under the legislative authority of the California Health and Safety Code § 2040. The field studies did not involve endangered or protected species.

Collected mosquitoes were identified to species using a dissection microscope and chill table (BioQuip, Rancho Dominguez, CA). Individual whole mosquitoes were placed into 2 ml microcentrifuge bead mill tubes that contained 2.8 mm ceramic beads (Fisher Scientific, Waltham, MA) and frozen at -20°C until use. Susceptible *Cx. tarsalis* utilized in insecticide CDC bottle bioassays described below were from the Kern National Wildlife Refuge (KNWR) colony [16, 17] and resistant *Cx. tarsalis* maintained in an insectary that were originally collected during 2019 in Woodland, California USA (Conaway strain; GPS coordinates: 38.647287, -121.668173). These strains were also used in the *Culex* RT*kdr* assay as controls for susceptible (wildtype KNWR strain) or resistant (mutant Conaway strain) *Cx. tarsalis*.

### 2. RNA Extraction

Individual whole mosquitoes were homogenized in 200 µl of MagMAX Lysis/Binding Buffer that was diluted 1:2 in phosphate buffer saline for 45 s using a Fisherbrand Bead Mill 24 Homogenizer (Thermo Fisher Scientific, Waltham, MA). RNA was extracted using the MagMAX-96 Viral RNA Isolation Kit and KingFisher Duo Prime Purification System programed with the MagMAX Pathogen Standard Volume software protocol as described by the manufacturer (Thermo Fisher Scientific, Waltham, MA) with the following exceptions: 80 µl of homogenate was extracted, magnetic beads were washed with 250 µl of wash solution, and the RNA was eluted in 50 µl. Notably, we employed the same method to extract RNA from mosquitoes that is widely used when testing for the presence of arboviruses [18]. Alternatively, RNeasy Plus Mini Kits (Qiagen, Mississauga, Ontario, Canada) were used to extract RNA from mosquitoes, as recommended by the manufacturer (Qiagen, Mississauga, Ontario, Canada). RNA concentration in the samples was measured using a NanoDrop 2000 Spectrophotometer (ThermoFisher Scientific, Waltham, MA) according the manufacturer recommendations.

### 3. RNAseq of *Vgsc* gene

*Vgsc* sequences were recovered from the host fraction of a metatranscriptomic RNAseq dataset derived from total RNA extracted from *Cx. erythrothorax* (N = 44) and *Cx. tarsalis* (N = 26) single mosquitoes collected from across California [19]. Sample collection, total RNA extraction, and paired-end mNGS RNAseq from each of the single mosquito specimens that served as input data here are described elsewhere ([19]; Sequence related archive: https://www.ncbi.nlm.nih.gov/sra/PRJNA605178). Raw fastq R1 and R2 data from each mosquito were first compressed to a unique set of reads sharing < 95% sequence identity via CD-HIT software [20, 21]. Translated blastx alignment of the resulting R1 and R2 reads with a representative *Vgsc* protein sequence from *Culex quinquefasciatus* Say (NCBI protein accession AFW98419.1; [22] was applied to identify deduplicated R1 and R2 reads from each mosquito sample which showed >50% of their length aligned with >90% identity to the *Cx. Quinquefasciatus Vgsc* reference sequence. Seqtk software (https://github.com/lh3/seqtk) was used to compile the separate *Culex erythrothorax* and *Cx. tarsalis* fastq reads that met these criteria from the 44 *Cx. erythrothorax* or 26 *Cx. tarsalis* individually deduplicated datasets. Partners of unpaired reads included in each pool were identified and included to ensure a full complement of paired reads, including additional potentially divergent *Vgsc* sequences that were not captured in the alignment step.

A total of 410 *Culex tarsalis* input read pairs and 481 *Culex erythrothorax* read pairs were carried forward from this step. Trimmomatic software [23] was used to remove the Sequencing library adapter sequences, along with low quality terminal bases of the reads. The resulting paired-end pooled datasets were each then separately used as input for SPAdes [24] paired-end *de novo* assembly of *Vgsc* transcripts. To facilitate *Vgsc* contig coverage analysis, read pools were aligned back to each of the identified *Vgsc* contigs via Bowtie2 [25].

The *Cx. tarsalis* and *Cx. erythrothorax* contig assemblies were aligned to the NCBI nt and nr databases via blastn and blastx, respectively, to identify the set of *de novo* assembled contigs that corresponded to *Vgsc* transcripts. A single 6878 bp *Cx. tarsalis* contig and two 6364 bp and 506 bp *Cx. erythrothorax* contigs were identified for further analyses. The *Cx. tarsalis* 6878bp contig encompasses an uninterrupted 2113 amino acid open reading frame, with additional 5’ 321 bp and 3’ 218 bp flanking terminal sequences.

The most closely related sequences in NCBI to this contig corresponded to several *Culex* complete *Vgsc* nucleotide and protein coding sequences. The best match was the *Cx. pipiens pallens* strain SS sodium channel mRNA (NCBI accession numbers KY171978.1 and ARO72116.1), showing >95% overall sequence identity at both the nucleotide and amino acid level. The *Cx. erythrothorax* contigs were not joined in the initial de novo assembly; however, the blastn and blastx alignment termini indicated a short (< 10 bp) region of overlapping sequence at the ends of these 2 contigs. Manual joining of these 2 contigs generated a 6709 bp contig that encodes an uninterrupted open reading frame of 2109 amino acids, and additional flanking 283 bp of 5’utr and 99 bp of 3’utr sequences. The best match is the *Cx. quinquefasciatus* isolate S-Lab sodium channel mRNA, complete cds (NCBI accession numbers EU817515.1 and ARO72116.1), showing >95% overall sequence identity at both the nucleotide and amino acid level. Accession numbers for the recovered *Cx. erythrothorax* and *Cx. tarsalis Vgsc* transcript sequences are MW176091 and MW176090, respectively. Resulting assemblies were manually reviewed via Geneious software (version 2019.0.4; https://www.geneious.com/) to generate final contig consensus sequences.

### 4. Detection of kdr SNP by RT-PCR

The primer and probe sequences to detect the *kdr* SNP were designed using Primer3Plus software (Table 1; [26]) based on the cDNA of *Vgsc* from *Cx. tarsalis* and *Cx. erythrothorax* (GenBank No. MW176090 and MW176091, respectively). Wildtype and mutant probes were labeled with fluorescein (FAM) and hexachlorofluorescein (HEX), respectively (Integrated DNA Technologies, Coralville, Iowa). A diagram depicting primer and probe locations, the 1014 mutation and intron site for *Vgsc* of *Cx. tarsalis* is provided in Figure 1. Nucleotide sequences were aligned using Basic Local Alignment Search Tool [27].

**Table 1.**
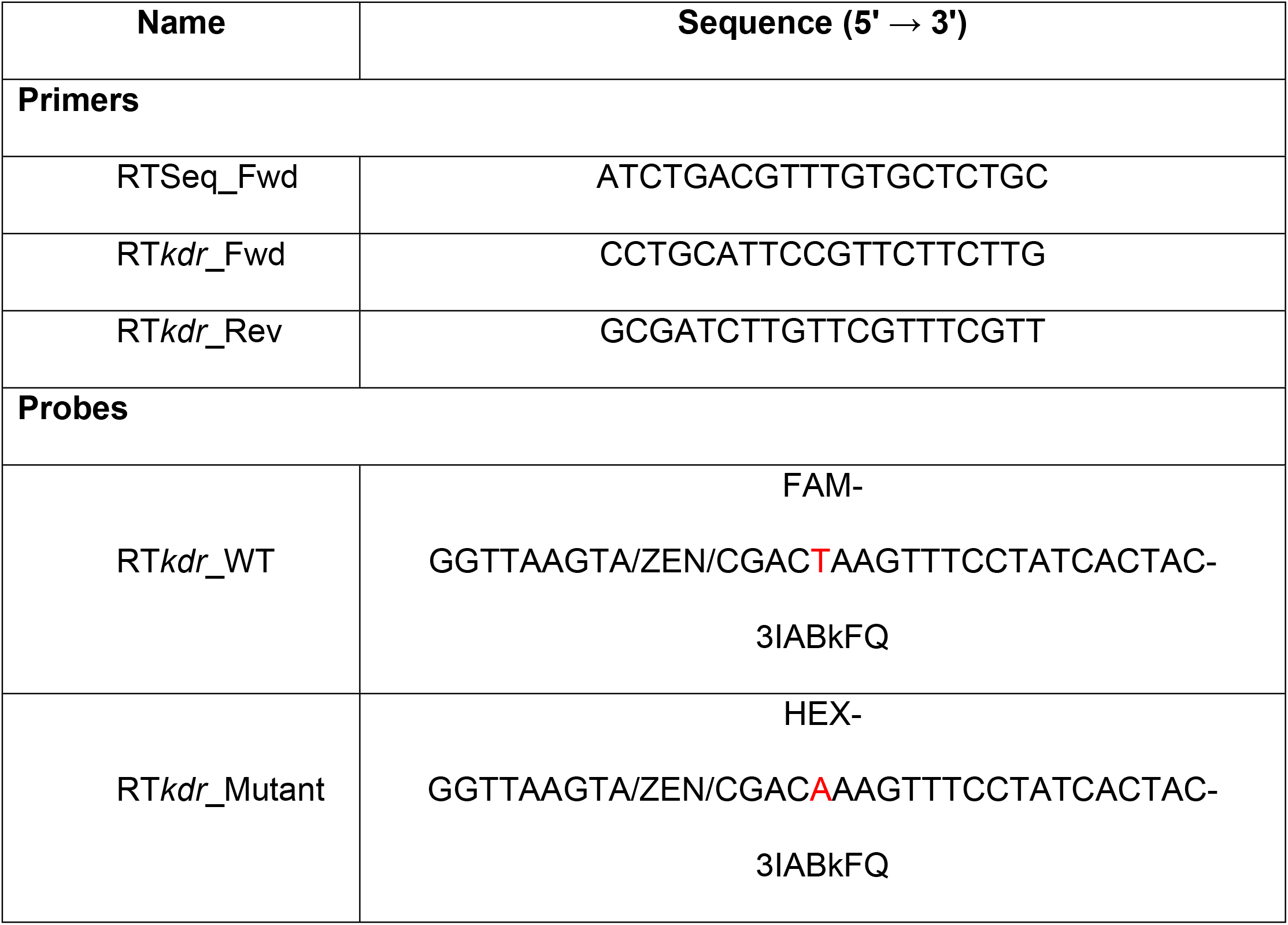
Primers and probes used in the *Culex* RT*kdr* assay. Red text indicates the location of the *kdr* SNP.

**Figure 1.**
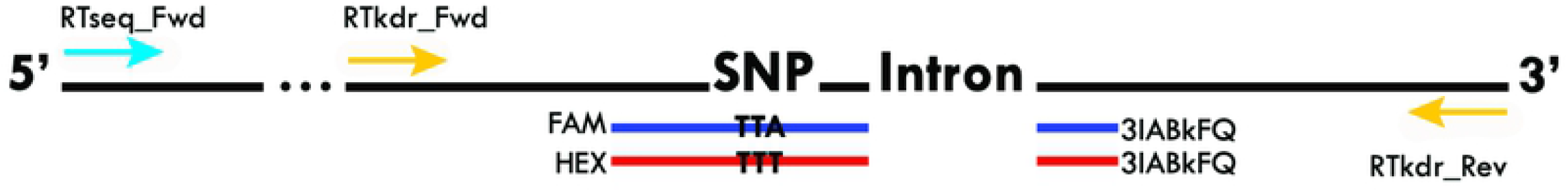
Schematic representation of sequencing primer (cyan), PCR primers (yellow), probes (red and blue), SNP and the intron of the kdr loci in *Vgsc* for *Cx. tarsalis*.

The Taqman Fast Virus 1-Step Master Mix (Thermo Fisher Scientific, Waltham, MA) was prepared as described by the manufacturer using 1 µl of template RNA (48.8-144.8 ng/µl), primers diluted to 900 nM and probes diluted to 250 nM. PCR plates were vortexed for 10 s at the highest setting (Fisherbrand™ Analog Vortex Mixer, ThermoFisher Scientific, Waltham, MA), centrifuged for 15 seconds (MPS 1000 Mini PCR Plate Spinner, Labnet International, Inc., Edison, NJ) and subsequently analyzed with a QuantStudio™ 5 Real-Time PCR System (Thermo Fisher Scientific, Waltham, MA) using the Genotyping setting which assigns sample results based on a proprietary algorithm. Amplification curves were reviewed manually to ensure algorithm accuracy. RT-qPCR cycling conditions were as follows: 50°C for 5 minutes, 95°C for 20 seconds, followed by 40 cycles of 95°C for 3 seconds and 60°C for 30 seconds. Primer and probe concentration and PCR cycling conditions were optimized to discriminate homozygous and heterozygous genotypes. Allele controls were added in the form of a no template control and a known susceptible control for *Cx. tarsalis, Cx. pipiens* and *Cx. erythrothorax*. A known resistant control was also included for each of the former except *Cx. erythrothorax* because a resistant specimen of that species was not found in the current study.

### 5. Validation Methods

#### 5.1 Insecticide Susceptibility Assays

CDC bottle bioassays were conducted to evaluate the resistance of adult mosquitoes to insecticides, according to CDC guidelines [13]. Three replicate bottles were evenly coated with 1 ml of technical grade insecticide (43 µg permethrin or 22 µg deltamethrin) that was diluted in acetone. Control bottles contained only acetone diluent. The diluent was evaporated from the bottles in the dark at room temperature. Adult female mosquitoes were transferred to the bottles (14-23 mosquitoes per bottle), and the number of dead or knocked down mosquitoes was recorded at 15 min intervals for 180 min. A mosquito was recorded as dead or knocked down if it could not stand unaided when the bottle was gently rotated; otherwise, the mosquito was counted as alive. Live and dead mosquitoes were separated, tested with the *Culex* RT*kdr* assay and the PCR products sequenced. Resistance ratios were calculated using the proportion of dead mosquitoes at the 45 min time point when average mortality was less than 100% with those from the susceptible Conway strain in the denominator.

#### 5.2 Validating the Culex RTkdr assay using Cx. pipiens quantitative PCR (qPCR) Taqman assay

The *Cx. pipiens* quantitative PCR (qPCR) Taqman assay that was developed previously [28] was used to validate the *Culex* RT*kdr* assay using *Cx. pipiens* samples. We followed the protocol for Taqman Multiplex Master Mix (ThermoFisher Scientific, Waltham, MA) with the following exceptions: BSA was excluded and nucleic acid that was isolated with the MagMAX-96 Viral RNA Isolation Kit (described above) was used as the template. We evaluated 75 *Cx. pipiens* mosquitoes using both the Taqman qPCR and *Culex* RT*kdr* assay and results were compared. Discordant samples were evaluated by sequencing the PCR products.

#### 5.3 Sequencing of PCR Products

PCR products were submitted to Elim Biopharmaceuticals (Hayward, CA) for PCR cleanup and sequencing. Because the RT*kdr*_Fwd primer is in close proximity to the SNP, we designed a sequencing primer further upstream in the *Cx. tarsalis* mRNA sequence that produced a 373 bp PCR product (RTseq_Fwd, Figure 1, Table 1). Primer, probe and template concentrations and PCR cycling conditions to generate PCR products for sequencing were as described above. Sequences were aligned to the *tarsalis Vgsc* mRNA sequence using MUSCLE [29] to locate the *kdr* SNP. Chromatograms were examined using 4Peaks software (Nucleobytes, Amsterdam, The Netherlands) to determine if heterozygosity was present at the SNP site.

### 6. Analyzing the Geographic Distribution of the kdr SNP

Tableau Software (Seattle, WA) was used to map the geographic distribution of the *kdr* SNP in mosquitoes that were collected in Alameda County (CA, USA). Allelic data for mosquitoes that were collected within 1 km of each other were combined. The trap sites were binned into two geographic regions, bayside and inland, that are separated by the San Francisco East Bay Hills, a natural boundary that limits movement of mosquitoes between the two regions. The distribution of allelic frequency was assessed by mosquito species and by geographical region (inland and bayside) within Alameda County. The resistant allele frequency, F_R_, in each population was estimated (Equation 1) where 2N_RR_ is the number of homozygous resistant mosquitoes, N_RS_ is the number of heterozygous resistant and N is the mosquito population size.

Equation 1. Equation for calculating resistance allele frequency

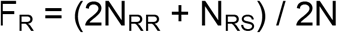

Associations between genotype, Y, and mosquito species, region of collection, and area type surrounding the collection site were estimated using Equation 2 from an ordinal logistic regression model with ordered outcome categories (SS, RS, RR). The model was fit using the polr function from the MASS [30] package in R Software (version 3.5.0;[31]) and used to estimate unadjusted and adjusted odds ratios for each variable. Figures were generated using ggplot2 software [32].

Equation 2. Equation for ordinal logistic regression model Logit

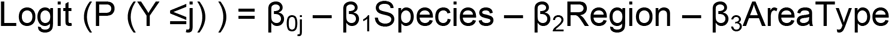

## Results & Discussion

### 1. Sequence Alignments

The *Vgsc* -1 cDNA sequences for *Cx. tarsalis* (GenBank No. MW176090), *Cx. erythrothorax* (GenBank No. MW176091) and *Cx. pipiens* (GenBank No. KY171978; [33]) were aligned using BLAST. There was 95.7% identity with the greatest divergence coming from the *Cx. pipiens* sequence. The forward and reverse primers matched 100% for all three species, however there were two mismatched nucleotides in the probe for *Cx. pipiens* (Figure 2, red boxes). These mismatches in *Cx. pipiens* resulted in less RT-PCR product relative to *Cx. tarsalis* and *Cx. erythrothorax* and no cross amplification of the wildtype and mutant RT-PCR probes as was observed for *Cx. tarsalis* and *Cx. erythrothorax* (Figure 3).

**Figure 2.**
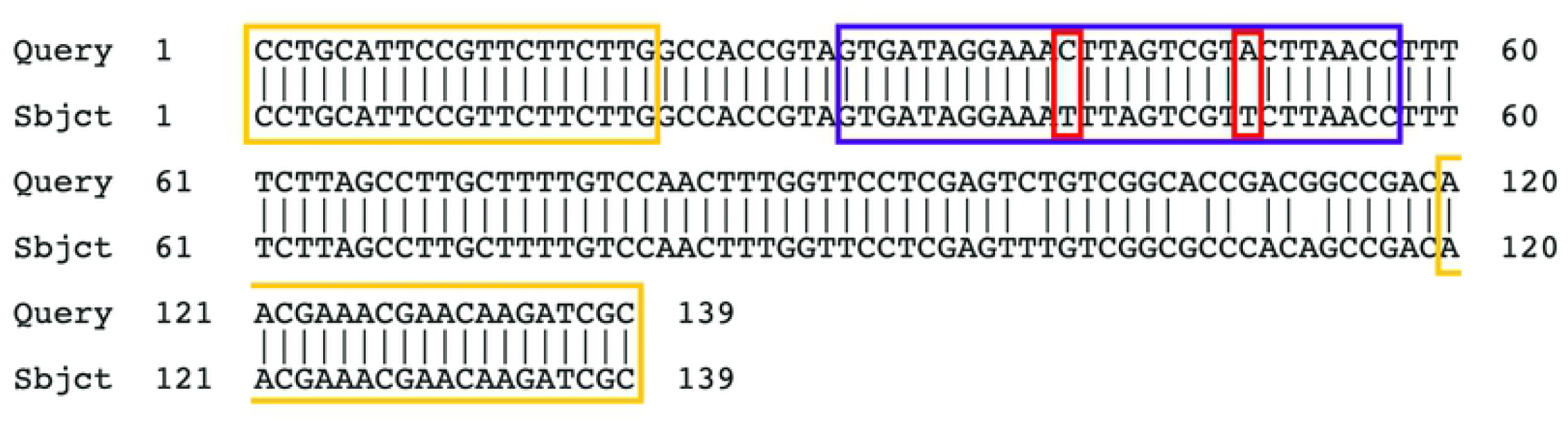
Basic Local Alignment comparing *Cx. tarsalis* (Query, Genbank No. MW176090) to *Cx. pipiens* (Subject, Genbank No. KY171978). Yellow boxes denote location of forward and reverse primers, purple box denotes probe location and red are mismatched bases.

**Figure 3.**
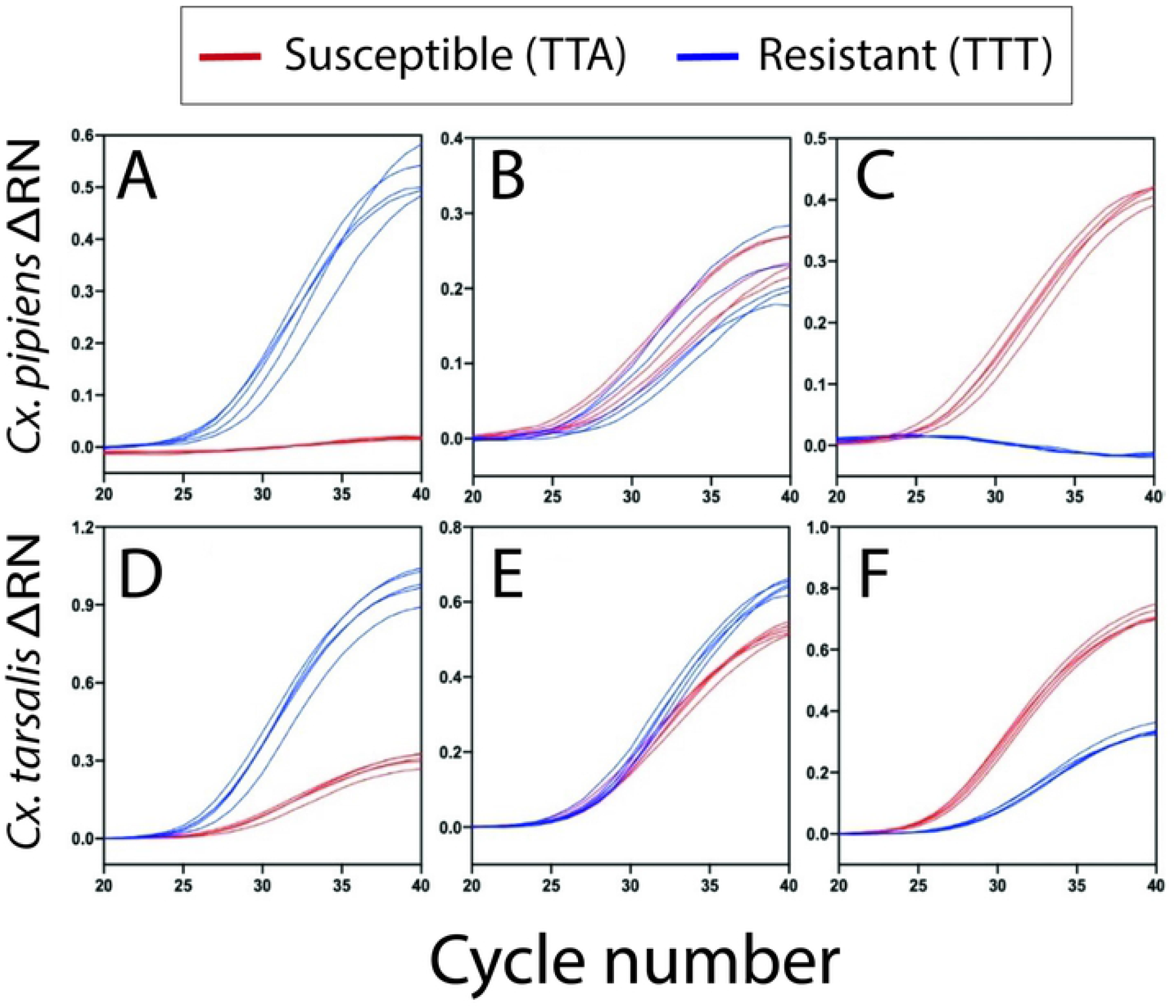
Amplification plots (ΔRN vs Cycle Number) with the wildtype probe labeled in blue and mutant probe in red. **(A)** *Culex pipiens* homozygous wildtype **(B)** *Culex pipiens* heterozygous **(C)** *Culex pipiens* homozygous mutant (**D)** *Culex tarsalis* homozygous wildtype (**E)** *Culex tarsalis* heterozygous **(F)** *Culex tarsalis* homozygous mutant.

### 2. Interpreting RT-PCR Results

Increased FAM or HEX fluorescence indicated a homozygous wildtype or mutant genotype, respectively (Figures 3A, 3C, 3D, 3F). A similar quantity of FAM and HEX fluorescence indicated that the specimen had a heterozygous genotype (Figures 3B, 3E). Allelic discrimination plots for *Cx. pipiens* and *Cx. tarsalis*, respectively, were used to identify outliers (Figure 4). Performance of the assay was also assessed using ΔCT values between the two probes as described below.

**Figure 4.**
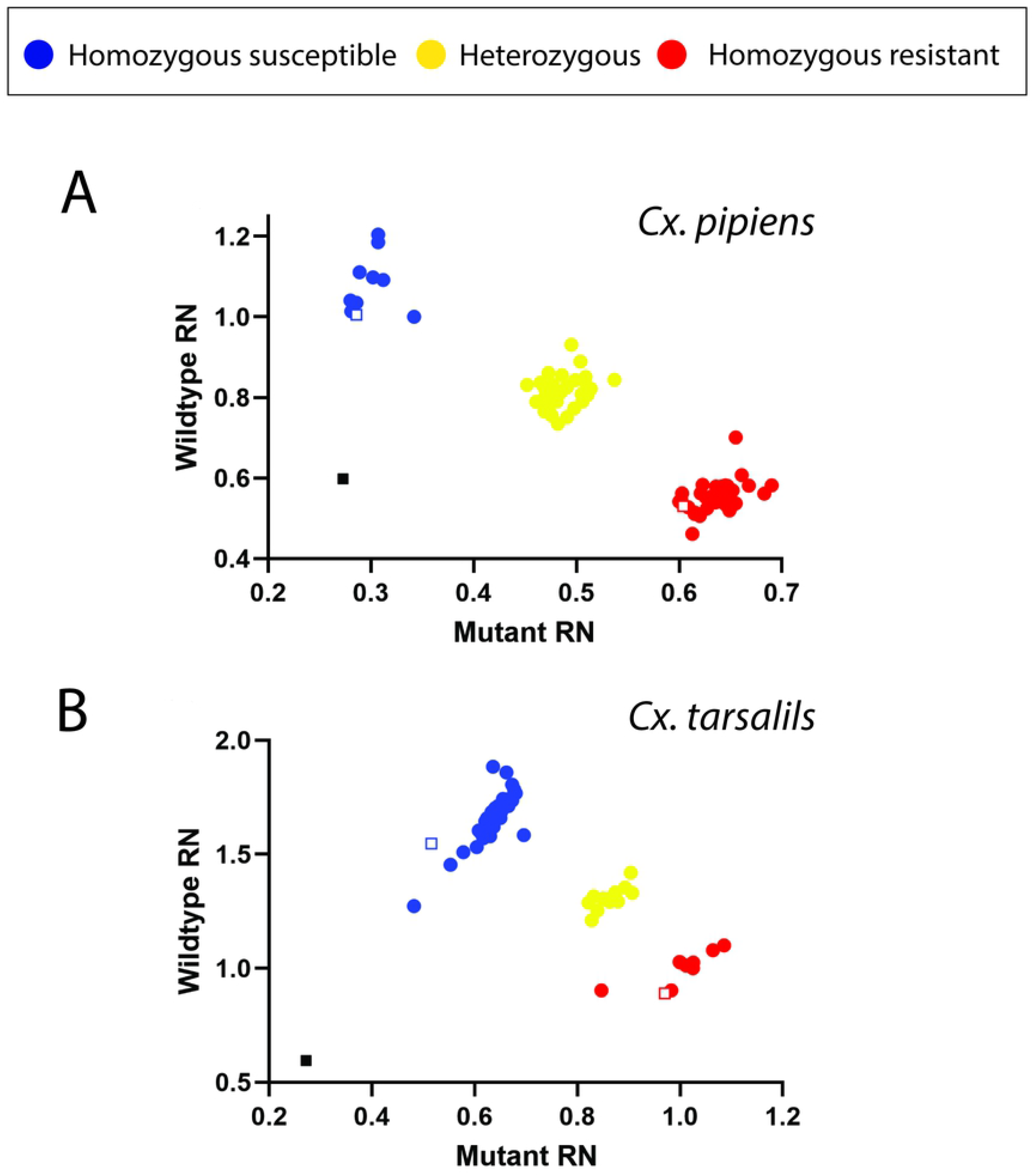
Allelic discrimination plots (Wildtype RN vs Mutant RN) for **(A)** *Culex pipiens* and **(B)** *Culex tarsalis*. Homozygous mutants are labeled with red ellipses, heterozygous with yellow ellipses, and homozygous wildtype are labeled with blue ellipses. Homozygous controls are labeled as open squares, outlined in their respective color. No template controls are labeled as black squares.

The ΔCT values for both *Cx. tarsalis* and *Cx. pipiens* were analyzed to determine a cutoff value for homozygous and heterozygous samples. For *Cx. tarsalis* the average ΔCT value of the heterozygous genotype was 0.264 ± 0.238 with a range of 0.008 – 0.932. Based on this information, *Cx. tarsalis* samples with a ΔCT of <1 were considered heterozygous. The average ΔCT values for mutant homozygous *Cx. tarsalis* was 2.944 ± 0.413 with a range of 2.077 – 3.964 and an average of 3.107 ± 0.782 with a range of 2.044 – 6.593 for homozygous wildtype. Therefore, samples with ΔCT values >2 were considered homozygous. ΔCT values for *Cx. erythrothorax* resembled *Cx. tarsalis* for homozygous wildtype, and was the only genotype detected for that species. Heterozygous ΔCT values were used to determine the *Cx. pipiens* genotypes because the opposing probes did not typically cross amplify for homozygous samples. Rarely, the opposing probe in *Cx. pipiens* samples amplified to produce a large ΔCT value. The average ΔCT of heterozygous *Cx. pipiens* samples was 0.981 ± 0.396 with a range of 0.164 – 1.779. Based on these findings, *Cx. pipiens* samples with ΔCT values under 2 were considered heterozygous and samples with undetermined or extremely large ΔCT values were considered homozygous.

Atypical amplification curves were occasionally observed for *Cx. pipiens* samples (< 5% of total), suggesting these mosquitoes may have been misidentified and were instead *Cx. erythrothorax. Culex pipiens* and *Cx. erythrothorax* are morphologically similar and can be mistaken for each other [4]. To help determine if the *Cx. pipiens* with uncharacteristic amplification curves may have been misidentified, we tested them using the *Cx. pipiens* qPCR assay that only produces a PCR product using nucleic acid isolated from *Cx. pipiens* or *Cx. quinquefasciatus* [28]. Each of those samples failed to amplify a product on the *Cx. pipiens* qPCR assay, providing additional evidence that the mosquitoes may have indeed been *Cx. erythrothorax. Culex tarsalis, Cx. pipiens, and Cx. erythrothorax* were the most prevalent *Culex* species collected during the study period. We also tested *Culex stigmatosoma* Dyar and *Culex apicalis* Adams. The low sample size for these species did not allow us to generate average ΔCT values. However, the amplification curves and sequenced PCR products were similar to *Cx. pipiens* or *Cx. tarsalis* (Figure S1), suggesting the *Culex* RT*kdr* assay may be effective for those species as well.

### 3. Validation Results

#### 3.1 Insecticide Susceptibility Assays

CDC bottle bioassays were used to identify *Cx. tarsalis* that were resistant or susceptible to permethrin and deltamethrin. Two lab-reared strains of *Cx. tarsalis* were assessed; one with known sensitivity to pyrethroids (KNWR strain) and another that displayed resistance (Conaway strain). The *Culex* RT*kdr* assay was used subsequently to determine the *kdr* SNP genotype of Conaway strain mosquitoes that were assessed in the bottle bioassay. Mortality or knockdown was on average less than 5% in mosquitoes placed in bottles that contained only diluent. At the 60 min time point, all susceptible strain (KNWR) mosquitoes had succumbed to permethrin and deltamethrin (Figure 5). At the 45 min time point, when the average mortality was less than 100% for all treatments, the Conaway strain was 54.5- and 58.8-fold more resistant to permethrin and deltamethrin, respectively. Resistance ratios of these magnitudes indicate that the Conaway strain was highly resistant to the insecticides. At 180 min, 21 ± 4% of the resistant Conaway strain mosquitoes had succumbed to permethrin and 38 ± 9% to deltamethrin (Figure 5). The slopes of the linear regression lines were significantly different for the Conway and KNWR strains (permethrin: F (1,50) = 309.2, P < 0.001; deltamethrin: F (1,50) = 50.84, P < 0.001), suggesting that their biological responses to the insecticides were different. The World Health Organization (WHO) classifies a population as resistant when mortality is below 90% [34]. The higher mortality rate observed in deltamethrin is expected as deltamethrin is a type II pyrethroid. Permethrin, a type I pyrethroid, may be more effective at knockdown because type I pyrethroids dissociate from the target faster than type II [12]. Because type II compounds, like deltamethrin, remain bound to the target longer, they are more effective at killing insects.

**Figure 5.**
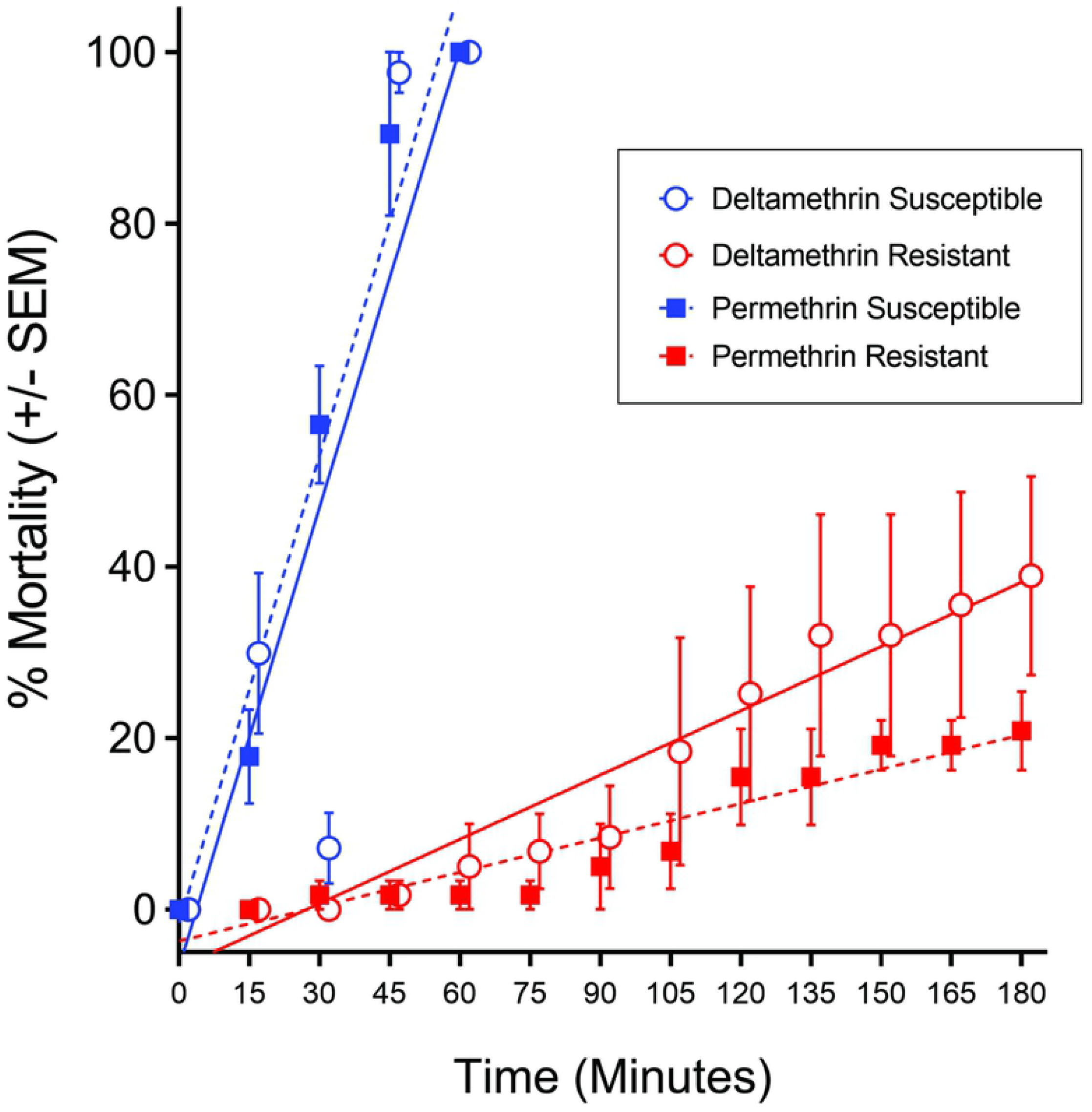
Bottle bioassay. Permethrin BBA results are depicted with closed squares with either blue or red representing permethrin susceptible or permethrin resistant, respectively. Deltamethrin BBA results are shown with open circles with either light blue or pink representing deltamethrin susceptible or deltamethrin resistant, respectively. Deltamethrin graphs are offset by 2 minutes for clarity. Equation of lines: Deltamethrin: susceptible KNWR strain, Y = 1.785*X - 6.627 (R^2^ = 0.7403); resistant Conaway strain, Y = 0.2499*X - 6.805 (R^2^ = 0.5130); Permethrin: susceptible KNWR strain, Y = 1.818*X - 1.553 (R^2^ = 0.9283); resistant Conaway strain, Y = 0.1337*X - 3.682 (R^2^ = 0.6443)

The genotype at the *kdr* SNP of the Conaway strain mosquitoes that survived or succumbed to permethrin or deltamethrin in the bottle bioassays was determined using the *Culex* RT*kdr* assay. All of the mosquitoes that survived exposure to permethrin or deltamethrin were homozygous mutant at the *kdr* SNP. Although the heterozygous, mutant and wildtype genotypes were observed only in mosquitoes that succumbed to the insecticides, there was no significant difference in the distribution of the genotypes in the bottle bioassays (Permethrin: F (2,2) = 18.21, P = 0.0521; Deltamethrin: F (2,2) = 5.569, P = 0.1522). Genotype results from the *Culex* RT*kdr* assay were confirmed by sequencing PCR products and viewing chromatograms. Chromatograms revealed a second *kdr* mutation at the 1014 amino acid among the resistant Conaway strain (Figure 6). Of the 47 BBA Conaway mosquitoes sequenced, 6 (13%) were heterozygous for the phenylalanine and serine *kdr* mutations. Both mutations were previously described in *Cx. quinquefasciatus* [35]. The serine *kdr* mutation may be associated with cross-resistance between DDT and pyrethroids [35]. Because DDT persists in the environment [36], it may have exerted a selective pressure on mosquitoes in the Conaway rice field that contributed to propagating the serine *kdr* mutation.

**Figure 6.**
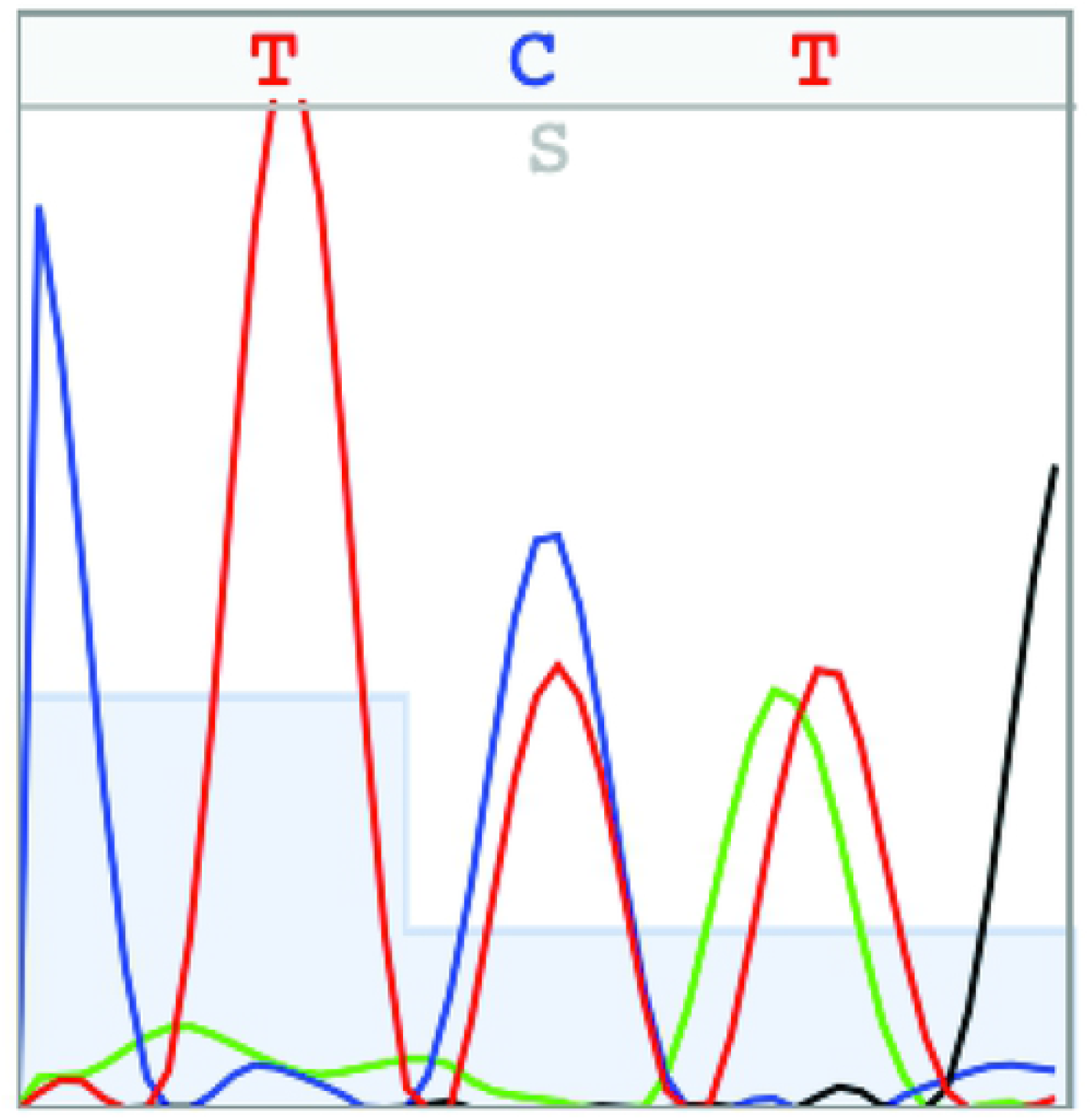
Chromatogram depicting heterozygosity for both the phenylalanine and serine/*kdr* mutations at the 1014 amino acid.

#### 3.2 Culex pipiens kdr detection Taqman qPCR

To determine the fidelity of the *Culex* RT*kdr* assay, individual *Cx. pipiens* mosquitoes were evaluated with both the *Culex* RT*kdr* and the *Cx. pipiens* qPCR assays (N = 75 mosquitoes). Three specimens (4%) failed to amplify a product after 30 PCR cycles in the *Culex* RT*kdr* assay and were excluded. Of the remaining mosquitoes, 69/72 (96%) results were concordant across both assays. Discordant results were sequenced to determine the correct genotypic call. Chromatograms for the three (4%) discordant samples indicated the mosquitoes were heterozygous and in agreement with the *Culex* RT*kdr* results, demonstrating that the *Culex* RT*kdr* assay was highly accurate (Table 2; paired t test, P > 0.9999).

**Table 2.** Validating the *Culex* RT*kdr* assay using a *Cx. pipiens* qPCR Taqman assay

| N | *Culex* RT*kdr* Assay |  |  | *Cx. pipiens* qPCR Assay |  |  |
| --- | --- | --- | --- | --- | --- | --- |
|  | TTA | TTA/T | TTT | TTA | TTA/T | TTT |
|  | (SS) | (RS) | (RR) | (SS) | (RS) | (RR) |
| 72 | 23 | 26 | 23 | 23 | 23 | 26 |

Susceptible homozygous (SS) TTA SNP, Heterozygous (RS) TTA/T SNP, Resistant homozygous (RR) TTT SNP

#### 4.3 Sequencing

PCR products from the *Culex* RT*kdr* assay were sequenced to further assess assay fidelity across five *Culex* species (N = 170; Table 3). Greater than 99% (169 out of 170) of the specimens were concordant with the sequencing and *Culex* RT*kdr* assay results (Table 3). The single discordant sample was misidentified as homozygous mutant by the *Culex* RT*kdr* assay, but the chromatogram revealed two peaks at the SNP location, indicating the mosquito was heterozygous (not shown). Using the sequencing results as the “true” result, we found the accuracy of the *Culex* RT*kdr* assay to be greater than 99%. High accuracy is common among both qPCR and qRT-PCR assays [37, 38]. Among the mosquitoes that were collected in Alameda County, only the L1014F mutation was found.

**Table 3.** Validating the *Culex* RT*kdr* assay by sequencing the resulting PCR products.

| Species (N) | RT-PCR |  |  | Sequencing |  |  |
| --- | --- | --- | --- | --- | --- | --- |
|  | TTA | TTA/T | TTT | TTA | TTA/T | TTT |
|  | (SS) | (RS) | (RR) | (SS) | (RS) | (RR) |
| *Cx. pipiens* (51) | 17 | 13 | 21 | 17 | 13 | 21 |
| *Cx. tarsalis* (97) | 18 | 17 | 62 | 18 | 18 | 61 |
| *Cx. erythrothorax*<br>(16) | 16 | 0 | 0 | 16 | 0 | 0 |
| *Cx. stigmatosoma*<br>(5) | 5 | 0 | 0 | 5 | 0 | 0 |
| *Cx. apicalis* (1) | 0 | 0 | 1 | 0 | 0 | 1 |
| Total (170) | 56 | 30 | 84 | 56 | 31 | 83 |

### 5. Distribution of Pyrethroid Resistance

The *Culex* RT*kdr* assay was used to assess the geographic distribution of the L1014F *kdr* mutation in Alameda County (Figure 7). Among the individual *Culex spp*. that were tested, 26.2% were homozygous resistant, 20.6% were heterozygous, and 53.3% were homozygous susceptible (N = 1383 mosquitoes). Ordinal logistic regression was used to determine associations between genotype, mosquito species, region of collection and area type. Because no resistance was identified in *Cx. erythrothorax*, ordinal logistic regression models were fit only to *Cx. pipiens* and *Cx. tarsalis* data. Table 4 summarizes the statistical results examining allele frequency and odds of resistant genotypes (heterozygous and homozygous resistant). The overall resistant allele frequency (F_R_) was highest among *Cx. pipiens* (0.57), low for *Cx. tarsalis* (0.15) and not present for *Cx. erythrothorax* (0.00). *Culex pipiens* had 8.99 times greater odds of being heterozygous or homozygous resistant compared to *Cx. tarsalis* (Table 4, 95%CI: 6.96 - 11.69). Adjusting for region and area type increased the association between resistance and *Cx. pipiens* (Table 4; OR: 11.01 (8.36 - 14.63)). The inland region revealed a higher F_R_ compared to the bayside region for both *Cx pipiens* and *Cx. tarsalis* (Figure 8). *Culex erythrothorax* was not present the inland region during the study period and all bayside *Cx. erythrothorax* were homozygous susceptible.

**Table 4.**
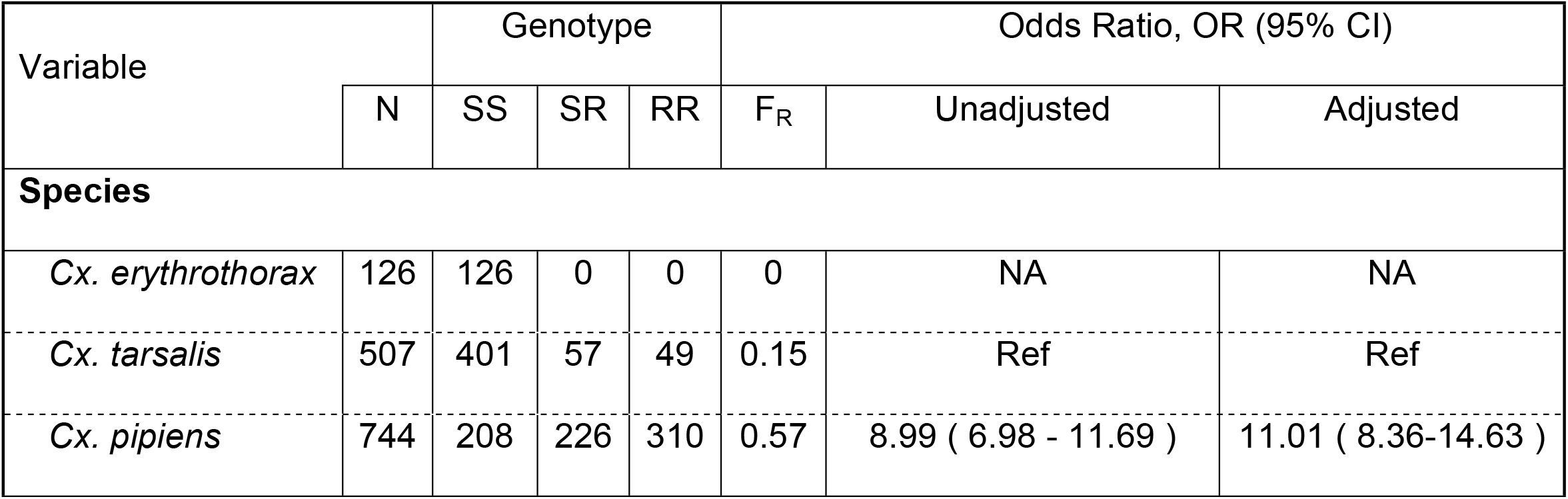
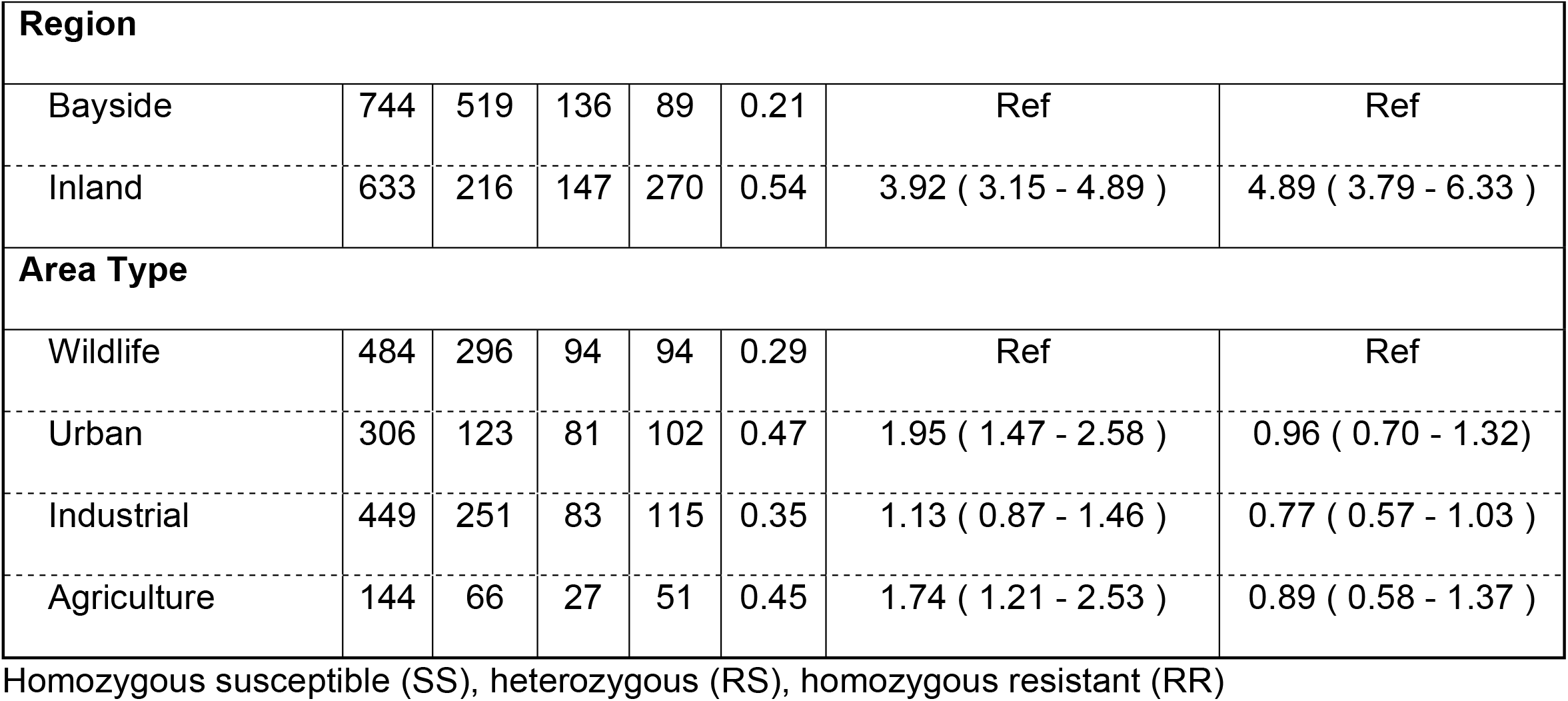
Genotypes detected, F_R_, unadjusted and adjusted odds rations among species, geographical region and land area type.

**Figure 7.**
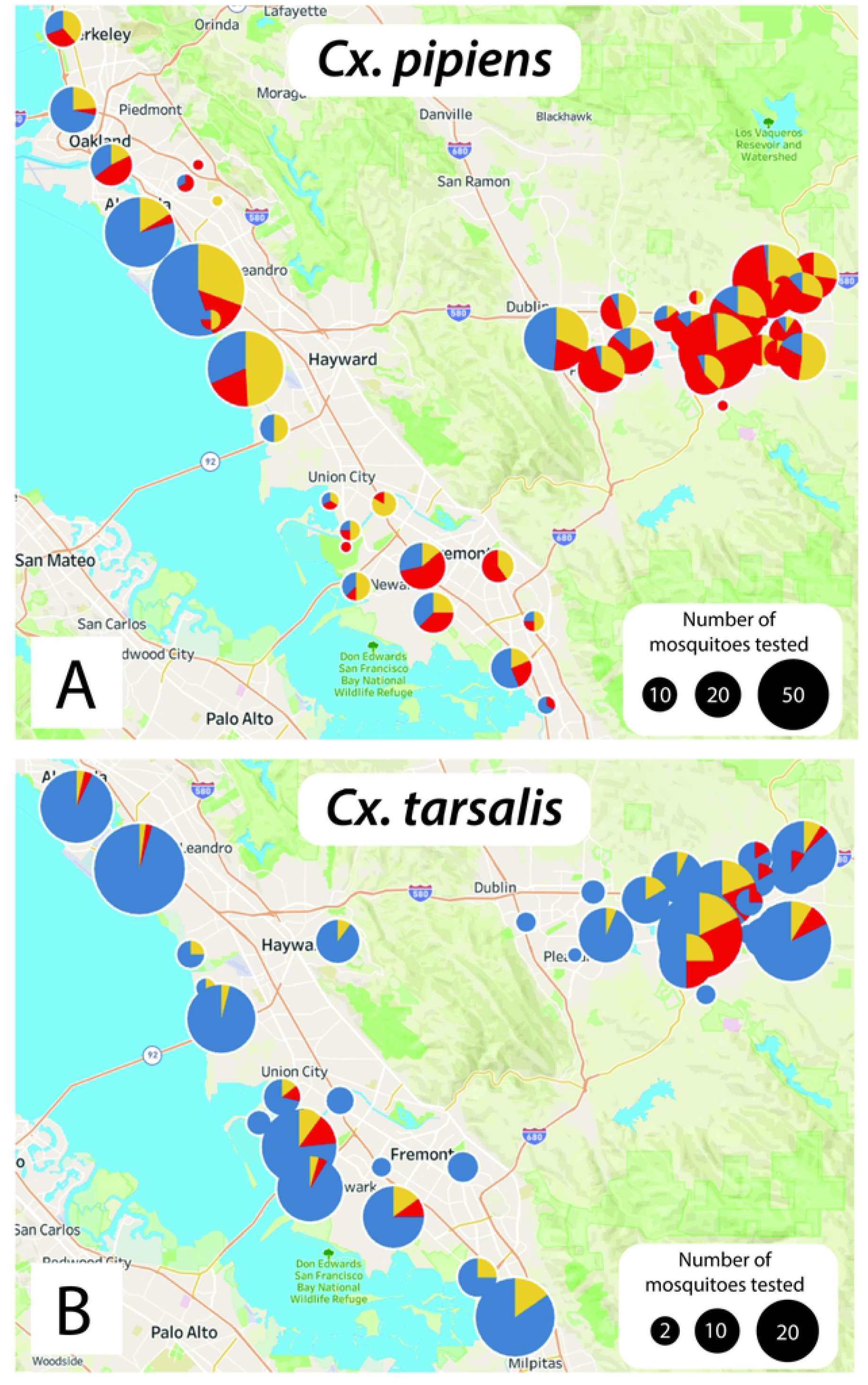
Resistance mapping within Alameda County showing **(A)** *Culex pipiens* and **(B)** *Culex tarsalis* in bayside and inland regions.

**Figure 8.**
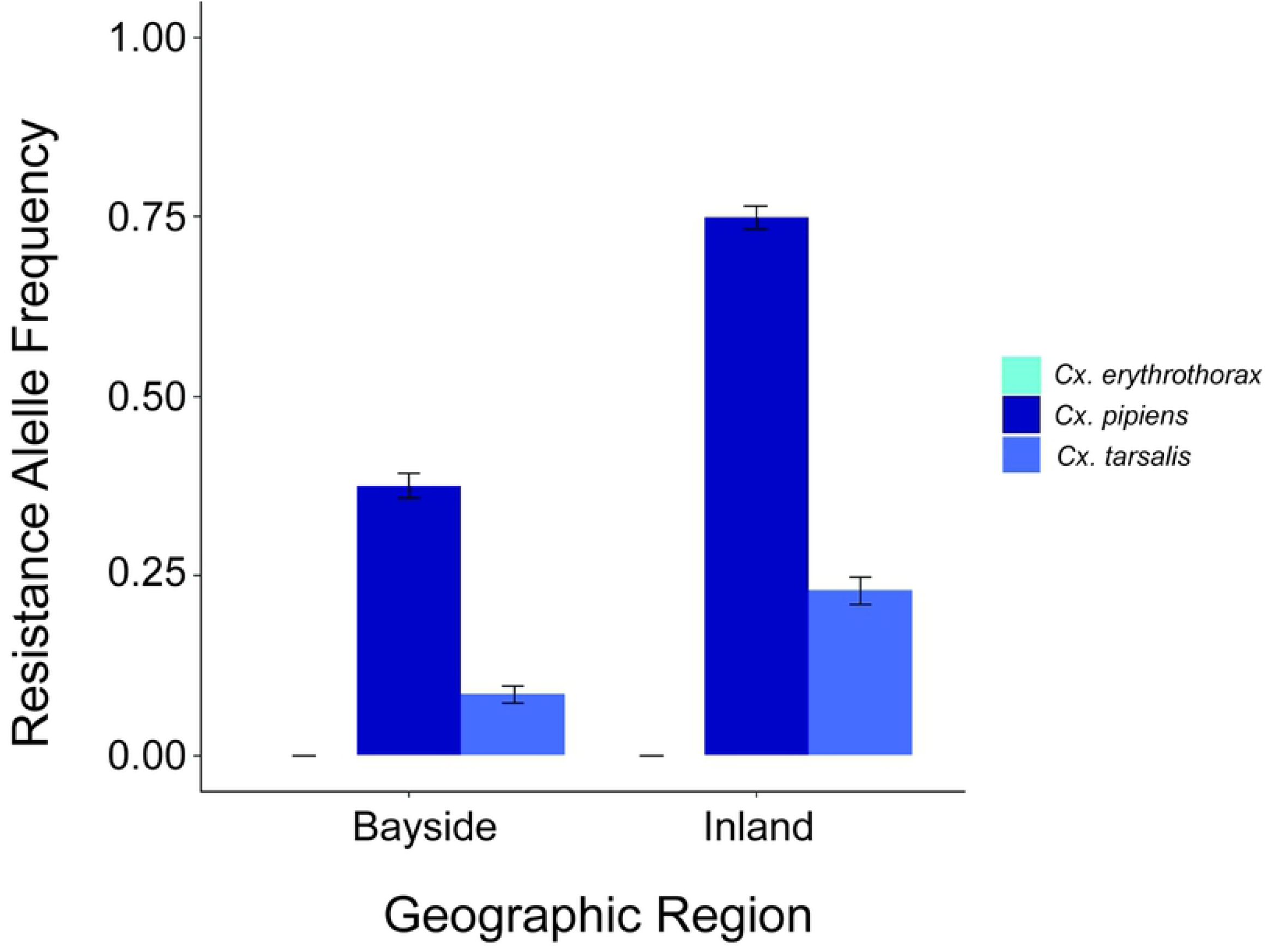
Resistant allele frequency (F_R_) of the L1014F *kdr* mutation by species and region. Bright blue, dark blue and medium blue bars represent F_R_ for *Cx. erythrothorax* (no resistance detected), *Cx. pipiens* and *Cx. tarsalis*, respectively. The F_R_ for bayside *Cx. pipiens* and *Cx. tarsalis* was 0.375 ± 0.018 and 0.0840 ± 0.012, respectively. The F_R_ for inland *Cx pipiens* and *Cx. tarsalis* was 0.749 ± 0.016 and 0.230 ± 0.016, respectively.

High resistant allelic frequencies were found previously in *Cx. pipiens* complex mosquitoes [11] [39]. *Culex erythrothorax* reproduce in heavily vegetated regions of shallow ponds and can be highly abundant in marsh habitats [40, 41]. While *Cx. erythrothorax* were typically found in bayside wetlands, *Cx. pipiens* and *Cx. tarsalis* were both present inland, yet the *Cx. pipiens* were far more resistant. Previous studies of urban creeks and outfall of storm drains in California found high levels of pyrethroids. The pyrethroids were proposed to have originated from homeowner use or structural pest control [20, 42]. Sites that were not near agriculture occasionally contained comparable levels of pyrethroids to those found in creeks near agricultural sites. The high levels of pyrethroids found in the sediments of the outfalls of storm drains may contribute to the resistance that we observed in *Cx. pipiens* as they reproduce in and around storm drain and storm drain outfalls [5, 43].

Mosquitoes from inland regions of Alameda County had elevated odds of containing the *kdr* SNP that is associated with pyrethroid resistance (Table 4; OR: 3.92 (3.15 - 4.89)). Adjusting for species and area type increased the association between resistance and mosquitoes that were collected from inland sites (OR: 4.89 (3.79 - 6.33)), suggesting an association present between inland mosquitoes and higher levels of resistance. The California Pesticide Information Portal (CPIP) shows that the top uses of pyrethroids in Alameda County were for structural pest control, wine grapes, almonds, pistachios and brussels sprouts (https://calpip.cdpr.ca.gov/main.cfm). While CPIP does not specify the township for structural pest control, using CPIP in conjunction with Pesticide Use Report (PUR) data, we were able to narrow agricultural pesticide use down to several locations within the inland region of Alameda County. Agriculture is widely practiced within the inland region of Alameda County and is less common in the bayside region. Studies of *Anopheles gambiae* suggest that insecticides from agriculture likely contribute to resistance in *Anopheles gambiae*, the malaria mosquito [21, 44, 45]. A similar pattern of pyrethroid use in agriculture cooccurring with pyrethroid resistance was observed in *Cx. pipiens* and *Cx. tarsalis*, two important vectors of WNV in North America.

## Conclusion

We developed a simple to use RT-qPCR assay that detects the *kdr* SNP that is associated with resistance to pyrethroid insecticides in at least five *Culex spp*. of mosquito. Like all PCR-based assays, the *Culex* RT*kdr* assay is not without limitations. It assay does not detect the serine *kdr* mutation that was discovered by sequencing the RT*kdr* assay PCR product from the Conaway strain (Figure 6). The serine *kdr* mutation suggests prior selective pressures, possibly from historical applications of pyrethroids or DDT. The *Culex* RT*kdr* assay also does not account for other pyrethroid resistance mechanisms such as overexpression or mutation of CYP9M10. Overexpression of CYP9M10 allows for increased detoxification of pyrethroids by cytochrome P450s monooxygenases [5, 46]. It was extensively validated for only *Cx. pipiens*, Cx. *tarsalis* and *Cx. erythrothorax* mosquitoes as we had a limited number of other *Culex* species available for the study. However, preliminary results suggest the assay performs for *Cx. apicalis* and *Cx. stigmatosoma*. Lastly, we know the assay performs well using Northern California mosquitoes, but genetic diversity across different countries may prevent the assay from detecting the L1014F mutation in these *Culex* species worldwide. More research is needed to determine whether this assay could be applied to mosquitoes collected outside of California.

Despite public health pesticide applications accounting for <1% of statewide pesticide use between 1993-2007 and with Alameda County Mosquito Abatement District having applied less than 300 ml of adulticide in the decade covering 2010 to 2020, pyrethroid resistance remains a concern [47]. Commercial use of insecticides for both structural and agricultural pest control may contribute to the higher pyrethroid resistance in mosquitoes from the inland region. In countries that ceased pyrethroid applications by vector control agencies, resistance remained high, likely due to household insecticides that contain pyrethroids [48].

The ability of the *Culex* RT*kdr* assay to perform well with multiple *Culex* species may benefit vector control agencies. It may be possible to apply this technique to other mosquito species as the V*gsc* sequences of *Aedes aeypti* Linnaeus and *Aedes albopictus* Skuse revealed a high percent identity around the V1016G *kdr* mutation, suggesting the development of an *Aedes* qRT-PCR assay may be possible [38]. Application of pyrethroids to a resistant population can potentially drive heterozygous populations (RS) to the homozygous resistant genotype (RR) further concentrates the and releases unnecessary chemicals into the environment. Prior to the development of this *Culex* RT*kdr*, there was no quantitative PCR assay to detect the L1014F mutation in *Cx. tarsalis*. The development of our *Culex* RT*kdr* assay satiates the need for a simple and reliable *Cx. tarsalis* PCR pyrethroid-resistance detection assay. We hope the assay will improve testing for pyrethroid resistance among *Culex* species.

## FIGURE LEGENDS

**Figure S1**. *Culex* RT*kdr* assay amplification curves for **(A)** *Cx. stigmatosoma* (N = 3, each was heterozygous) and **(B)** *Cx. apicalis* (N = 3, two were homozygous mutant, one was heterozygous,).

